# Inferring strain-level mutational drivers of phage-bacteria interaction phenotypes arising during coevolutionary dynamics

**DOI:** 10.1101/2024.01.08.574707

**Authors:** Adriana Lucia-Sanz, Shengyun Peng, Chung Yin (Joey) Leung, Animesh Gupta, Justin R. Meyer, Joshua S. Weitz

## Abstract

The enormous diversity of bacteriophages and their bacterial hosts presents a significant challenge to predict which phages infect a focal set of bacteria. Infection is largely determined by complementary – and largely uncharacterized – genetics of adsorption, injection, cell take-over and lysis. Here we present a machine learning approach to predict phage-bacteria interactions trained on genome sequences of and phenotypic interactions amongst 51 *Escherichia coli* strains and 45 phage *λ* strains that coevolved in laboratory conditions for 37 days. Leveraging multiple inference strategies and without *a priori* knowledge of driver mutations, this framework predicts both who infects whom and the quantitative levels of infections across a suite of 2,295 potential interactions. We found that the most effective approach inferred interaction phenotypes from independent contributions from phage and bacteria mutations, accurately predicting 86% of interactions while reducing the relative error in the estimated strength of the infection phenotype by 40%. Feature selection revealed key phage *λ* and *E. coli* mutations that have a significant influence on the outcome of phage-bacteria interactions, corroborating sites previously known to affect phage *λ* infections, as well as identifying mutations in genes of unknown function not previously shown to influence bacterial resistance. The method’s success in recapitulating strain-level infection outcomes arising during coevolutionary dynamics may also help inform generalized approaches for imputing genetic drivers of interaction phenotypes in complex communities of phage and bacteria.

## Introduction

Next-generation sequencing technology has revealed widespread diversity in microbial and viral communities (Aylward et al., 2017; Munson-McGee et al., 2018; Breitbart et al., 2018; Dominguez-Huerta et al., 2022; Sunagawa et al., 2015; Nayfach et al., 2021; Sunagawa et al., 2020). In parallel, the development of analytical tools to characterize species interaction networks from co-occurrence and/or time series data has led to a better understanding of microbial community structure and function (Faust and Raes 2012; Flannick et al., 2006; Stein et al., 2013; Berry and Widder 2014; Liao et al., 2020; Jiliang et al., 2022; Tamar and Kishony 2022). In principle, it should be possible to infer microbial interaction networks directly from genotypes and environmental context (Manrubia et al., 2021). In the case of phage-bacteria interactions, phage infection of a focal bacterial strain requires adsorption to specific cell-surface receptors (e.g., protein, lipid, carbohydrate) (Neurath et al., 1986; Wang, Hofnung and Charbit 2000; Chatterjee and Rothenberg 2012; Gaborieau et al. 2024), although in many cases the specific receptor remains unknown or modulated by poorly characterized biosynthetic pathways (Tetz and Tetz 2022). However, even if a phage adsorbs to a bacterium, there are many intracellular resistance mechanisms that could assist or inactivate phage infection altogether (Zborowsky and Lindell 2019; Koonin et al., 2020; Gao and Feng 2023). Categorizing effective, extracellular adsorption and intracellular replication remains challenging. Hence, despite significant progress in linking microbial genotype to phenotype, less progress has been made with understanding the genetics of traits that influence microbial species interactions (including virus and host pairs) given the additional complication that the phenotypic output of an association may depend on the joint effects of two separate genomes (Bajic et al., 2018; de Jonge et al., 2019; Buckling and Rainey 2002; Elena and Lenski 2003; Koskella and Brockhurst 2014; Poullain et al., 2008; Kaltz and Shykoff 2002; Beckett and Williams 2013; Weitz et al., 2013; Gurney et al., 2017).

The problem of understanding the genetic basis of interactions requires the development of computational approaches to construct genotype-to-phenotype maps. Conventional approaches try to correlate phenotypic differences with genetic variation (e.g., this is true for the broad scope of work in genome-wide associated studies (Horton et al., 2014; Falush 2016; Power et al., 2017)). The challenge for inferring interaction-associated phenotypes is that such interactions arise due to the combination of multiple genotypes (e.g., phage and host genotypes) leading to new combinatorial challenges. Initial steps towards interaction inference have been made through mutation-based association approaches that have successfully uncovered combinations of virus and host mutations that correlate with successful virus-host interactions (Tamar and Kishony 2022; MacPherson et al., 2018; Jallow et al., 2009; Scanlan et al., 2011; Borin, Lee, Lucia-Sanz et al., 2023; Boeckaerts et al., 2024). Conceptually, the challenge of uncovering interaction phenotypes is similar to attempts to tackle the problem of studying complex traits where gene-by-gene (G x G) interactions or gene-by-environment (G x E) interactions shape phenotypes (Wei et al., 2014; An et al., 2009; Gibson 2015; Gupta, Zaman et al., 2022).

In the case of virus-microbe systems, efforts to predict interaction phenotypes require leveraging specific system features and may depend on taxonomic scales. For example, computational approaches are increasingly used to predict the host range of viruses in a broad taxonomic sense, e.g., leveraging tetranucleotide frequencies and other sequence-specific information (Edwards et al., 2016; Dutilh et al., 2017; Roux et al., 2023, Bastien et al., 2024). However, predicting strain-specific interactions remains a difficult task, particularly because taxonomic markers are known to be a poor proxy for infection profiles (Sullivan et al., 2003; Kauffman et al., 2022). Recent studies have shown some improvement in resolving strain-specific interaction phenotypes, e.g., by using CRISPR spacers and metagenomic data to identify recent phage infection (Roux et al., 2021; Szabo et al., 2022; George and Hug 2023; Shang and Sun 2022) or by co-clustering phage and bacteria mutations, respectively, amongst strains that tend to interact as a means to identify associated gene or sequence differences (Kauffman et al., 2022).

Here, we link whole genome-wide changes in phage and bacteria with observed changes in interaction phenotypes in a coevolutionary context using a machine learning inference framework. We do so by leveraging emergent genotype and phenotype changes in coevolving populations of *Escherichia coli* B strain REL606 and bacteriophage *λ* strain cI26 during a 37-day experiment (Gupta, Peng et al., 2022). The key idea is to recapitulate infection phenotypes within an interaction network through a hierarchical regression approach without *a priori* assumptions about driver mutations or the nature of genetic interactions. In contrast, prior work on microevolutionary changes in infectivity have focused on changes to genes or proteins with known functions in model organisms (Gaborieau et al., 2024; Meyer et al., 2012; Lobo et al., 2009; Modi et al., 2013). Such approaches depend on existing annotation of genes or mutations, and thus are limited by both the quality and quantity of annotations. Our regression framework predicts a substantial portion of phage-host infection phenotypes, including: i) who infects whom and ii) with what efficiency. In doing so, we identify prioritized phage and bacterial mutations underlying changes in infection phenotypes and reveal that additive effects of phage and host mutations can be sufficient to predict interaction phenotypes. As we explain, this finding suggests a route to generate testable hypotheses for genome sites underlying interactions between phage and bacteria amongst closely related strains.

## Results

### Phage-bacteria mutation profiles and cross-infection matrix

We used genome sequences of 50 bacterial hosts (descended from E. *coli* B strain REL606) and 44 phage strains (descended from *λ* strain cI26) isolated at varying time points during a 37-day coevolution experiment (Gupta, Peng et al., 2022). Mutation profiles of the host and phage revealed many changes in their genomes, including 18 and 176 unique mutations for the host and phage, respectively (Supplementary Data S1 and Data S2). The interactions between phage and bacteria (including evolved strains and ancestors) were quantified via plaque assay in terms of the relative value of the efficiency of plating (EOP) in a target host by a focal phage strain compared to that in the sensitive ancestral host. Additional details of the EOP calculations are described in Methods section “Experimental setup and data collection” (Gupta, Peng et al., 2022). The interactions of all phage-bacterial pairs (including ancestors) yielded a 51 × 45 phage-bacteria cross-infection matrix. A summary of the mutation profiles and the phage-bacteria cross-infection matrix is shown in Fig. 1. We observed 913 successful (EOP > 0) and 1382 unsuccessful (EOP = 0) phage infections of 2295 phage-bacteria pairs. The distribution of EOP values was skewed, with 95% of values ranging from 0 to 1.5, including a long tail with a significant variability in the observed phenotypes (Supplementary Fig. S1). The co-occurrence of mutations in different genomic contexts suggested it might be feasible to infer host and phage mutations that disproportionately impact the phage-bacteria interaction phenotype (Supplementary Fig. S2).

**Figure 1.**
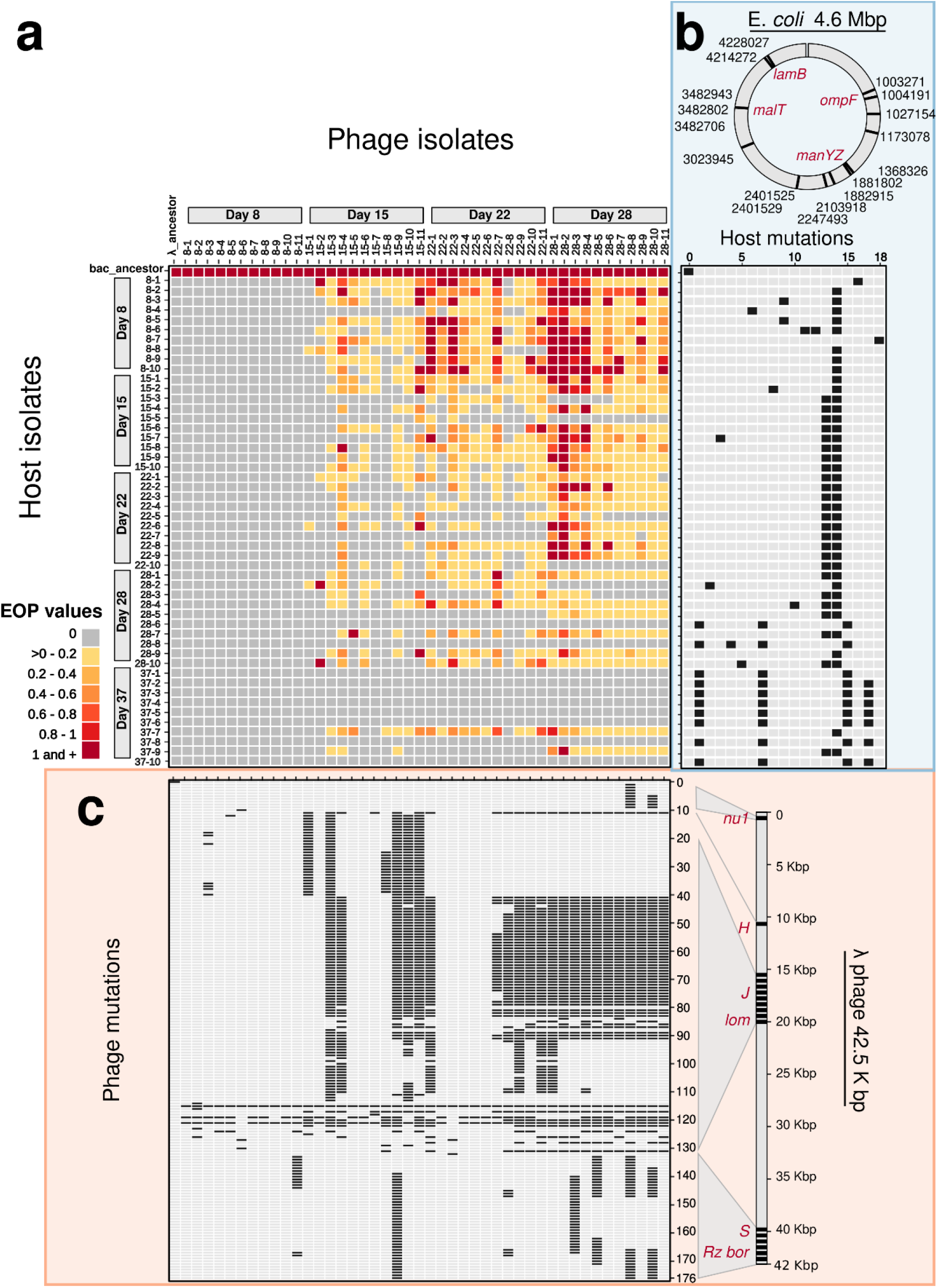
Phage-bacteria cross-infection matrix and mutation profiles. (a) Cross-infection matrix, including host and phage ancestor strains, and 50 bacteria (rows) and 44 phage (columns) strains isolated during 37-day coevolution experiment (day of isolation indicated). Names correspond to “day of isolation – number of isolate”. Colored cells are EOP values of infection as in legend, grey cells indicate no infection. (b-c) Mutation profiles for each isolate (positions mutated are in black and in grey otherwise) for 18 (host) and 127 (phage) found mutations numbered in sequential order of appearance in the corresponding genome. (b, in blue) Host isolates (rows) and mutation profiles (columns) for 1 to 18 unique mutations found in nt position 1,003,271 to 4,228,027 of the E. *coli* genome (c, in orange) Phage isolates (columns) and mutation profiles (rows) for 1 to 127 unique mutations found in nt position 175 to 42,491 of the *λ* phage genome. For the complete list of host and phage mutations see Supplementary Data S1 and Data S2. Important genes for phage-host interaction are highlighted in red and discussed in the main text.

### Model for predicting a coevolution-induced phage-bacteria interaction network

We developed a framework for predicting the combined effect of mutational profiles in phage and bacteria on interaction phenotypes. The underlying framework uses a logistic regression approach to determine infection outcomes – specifically predicting the presence or absence of infection (referred to here as POA)– for each phage-bacteria pair in the cross-infection network (see Methods section “Framework design”). Additionally, we examined how combining different mutation profiles (i.e., phage and host genotypes separately or in combination) improves POA inference. We tested five models in total. Three of these models use linear combinations of mutation features: one for host-only mutations (H model), one for phage-only mutations (P model), and one for the addition of phage and host mutations (linear model). The other two models use nonlinear combinations of phage and bacteria mutations with a first-order term (nonlinear model) and with a second-order term (mixed model). A comprehensive description of each model is provided in the Methods section “Feature construction”.

By comparing the performance of the models, we find that those with combinations of phage and bacteria mutational features predict the original POA phenotypes significantly better than the null model. In particular, we find that the linear model outperforms all the rest in the validation step (*P* < 9.44e-5) with a mean classification accuracy of ∼86% (Supplementary Fig. S3a). The predicted and experimentally measured POA phenotypes are shown in Fig. 2 along with the associated coefficients of the mutational features resulted from the linear, nonlinear, and mixed models. The value and sign of the inferred coefficients indicate the contributions that each mutational feature has on the predicted phenotype: positive coefficients increase the probability of infection, and the opposite is true for negative coefficients. Notably, we observe that bacterial mutations are more likely to have a negative effect (negative coefficients) due to the evolution of host resistance, whereas phage mutations tend to have a positive effect (positive coefficients), indicating selection for counter-defense traits that expand host range (see (Gupta, Peng et al., 2022)). We show that the additive effects of phage and host mutations alone can recapitulate the POA matrix without explicit inclusion of higher order interaction effects. We perform feature importance analysis of the linear model (detailed in the Methods section “Final predictions and feature importance analysis”) which reveals 5 host mutations and 32 phage mutations that have a positive effect on predicting the phage-host interaction network, compared with 7 host mutations and 15 phage mutations that have a negative effect (Supplementary Fig. S4a, Supplementary Data S3).

**Figure 2.**
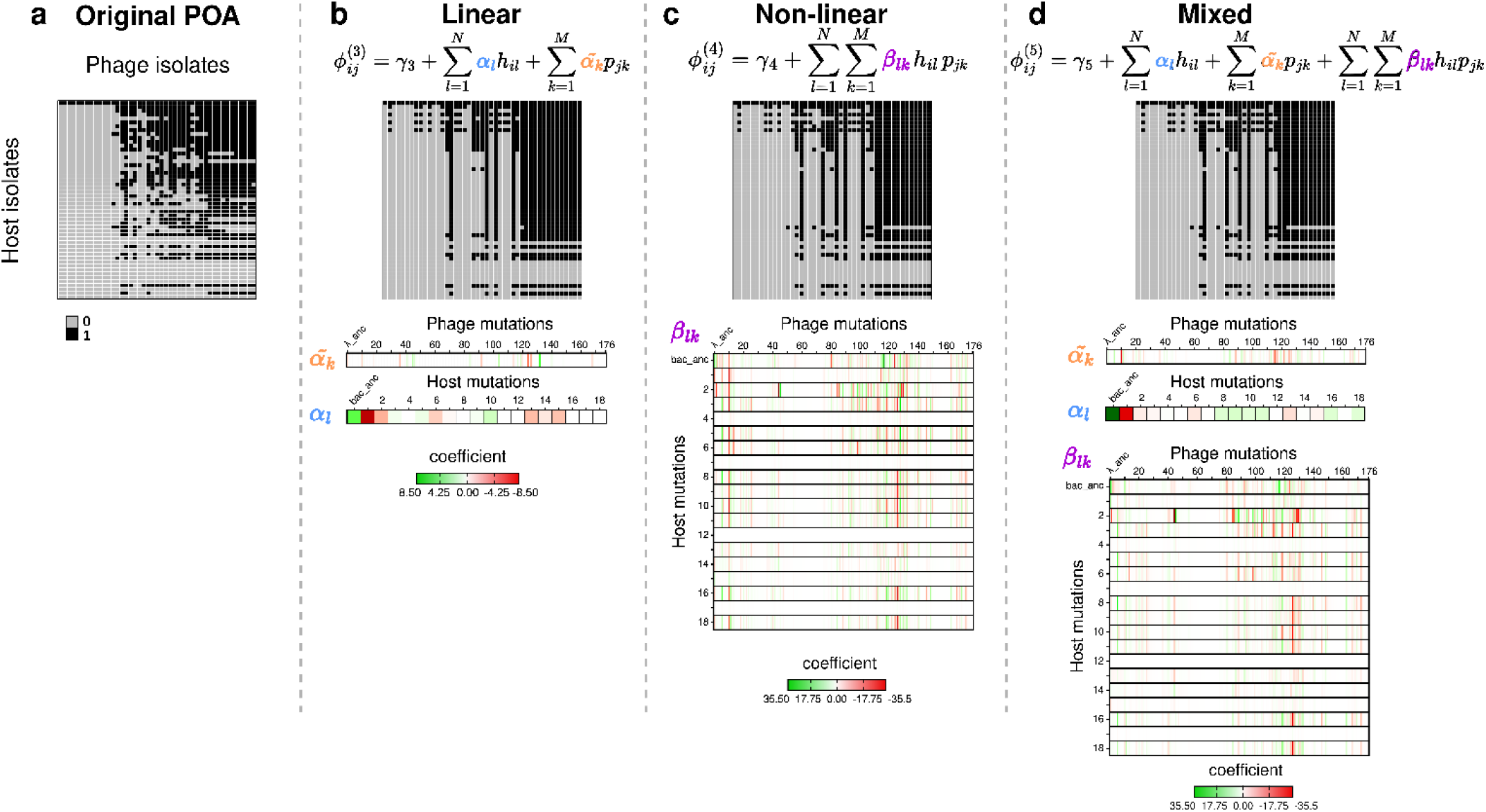
Model for predicting phage-host interaction network. (a) Original POA matrix showing presence (black) and absence (gray) of successful infection between phage (columns) and host (rows) isolated pairs. (b-d) Results of the different model predictions as of the POA matrices, and coefficient values for 176 phage and 18 host mutational features plus the ancestor trait using (b) a linear mutation set (equation [6]), (c) nonlinear mutation set (equation [8]) and (d) mixed combination of phage and host mutation set (for details see Methods section “feature construction”). The color of the coefficient indicates positive (green) to negative (red) effects of each mutational feature (phage: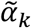, host: *α*_*l*_) or combination of mutations, *β*_*lk*_.

### Model for predicting the efficiency of phage-bacteria infections

Next, we extended the prediction framework to infer the efficiency of phage-bacteria interactions (EFF phenotype) as a function of phage and bacteria genotypes. EFF phenotypes were defined as log-transformed EOP measurements of individual infection pairs (see Methods section “Framework design” and Supplementary Fig. S5), while keeping the cross-interaction network fixed (Fig. 3a). Performances of the linear regression models based on the five different mutational feature sets (models H, P, linear, nonlinear, and mixed) were validated via mean absolute error (MAE). Results show that models containing combinations of phage and bacteria mutational features predict the original EFF phenotypes significantly better than the null model (Supplementary Fig. S3b; see panels in Fig. 3b-d). In particular, the linear model has the lowest validation MAE (*P* < 3.95e-14) with ∼40% reduction of the mean error compared to the null model. The coefficients associated to each mutational feature in the linear model denote the relative impact on the EFF phenotype. Feature importance analysis identified 8 host mutations and 25 phage mutations that increase the efficiency of phage infection (positive coefficients), compared to 6 host mutations and 28 phage mutations that reduce the efficiency of phage infection (negative coefficients). The full set of coefficients is listed in Supplementary Data S4, and driver mutations are shown in Supplementary Fig. S4a.

**Figure 3.**
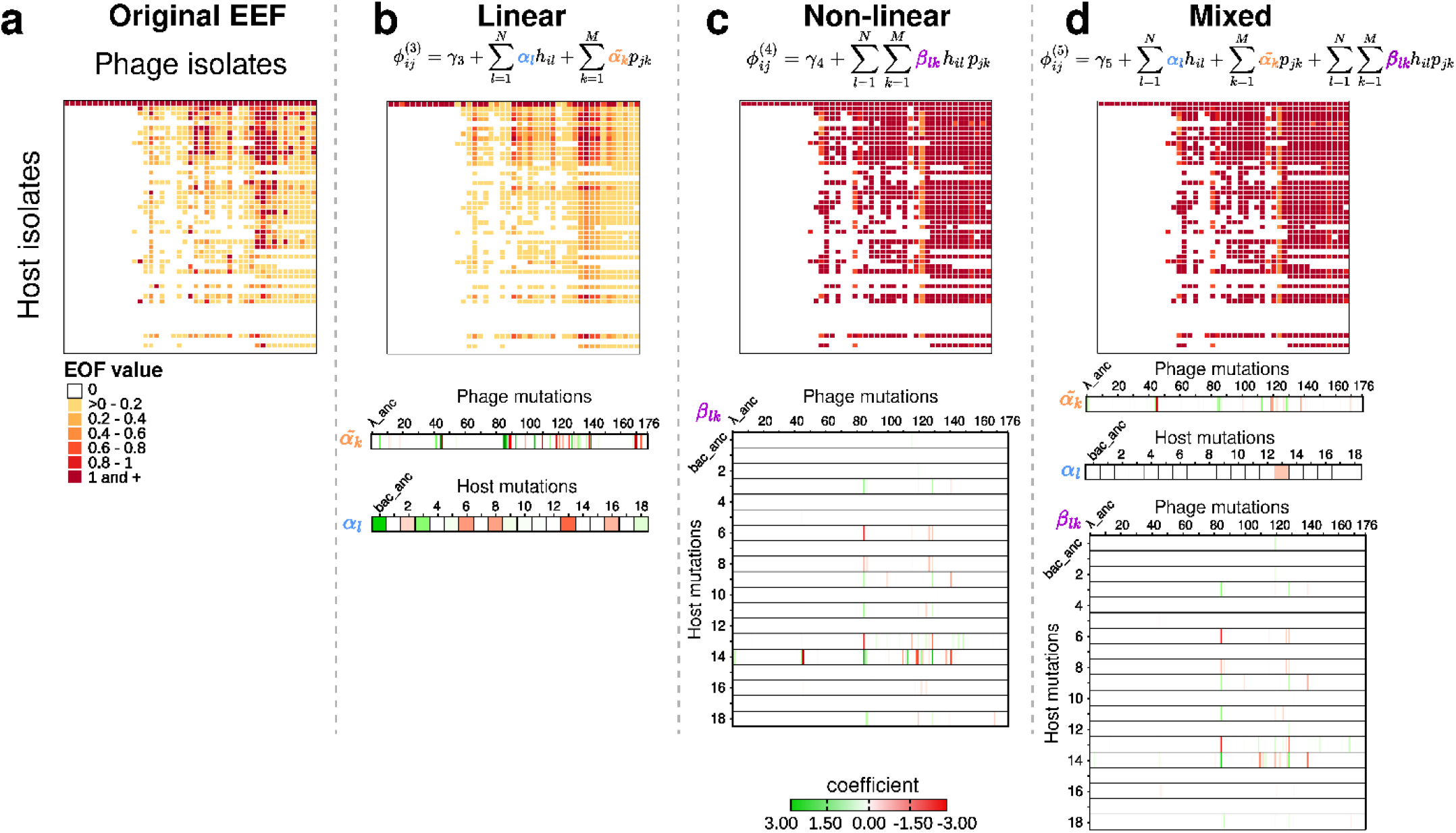
Model for predicting the efficiency of infection. (a) Original cross-infection matrix where colors are EOP values of infection between phage (columns) and host (rows) isolate pairs, white cells indicate no infection. (b-d) Results of the different model predictions as of the EFF matrices, and coefficient values for 176 phage and 18 host mutations plus the ancestor trait using (b) a linear mutation set (equation [6]), (c) nonlinear mutation set (equation [8]) and (d) mixed combination of phage and host mutation set (equation [10]) (for details see Methods section “feature construction”). The color of the coefficient indicates positive (green) to negative (red) effects of each mutational feature (phage: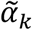, host: *α*_*l*_) combination of mutations, *β*_*lk*_.

### Characterizing the functional role of putative driver mutations of phage-bacteria interaction

The prediction framework yielded dozens of phage and bacterial mutations that significantly impact POA and EFF predictions (Fig. 2 and 3, respectively). In Fig. 4 we organize putatively important mutations revealed by the feature analysis using the linear models associated with POA (Fig. 4a, Supplementary Data S3) and EFF (Fig. 4b, Supplementary Data S4) matrices. We found 3 phage mutations and 1 bacterial mutation that show a significant positive effect for the POA phenotype. For phage, these mutations include 2 nonsynonymous mutations in genes *S* and *J* and a synonymous mutation in gene *J*. For the bacteria we identified a nonsynonymous mutation in the *ccmA* gene which encodes a subunit of an ABC transporter to the periplasm (Fig. 4a and Supplementary Fig. S4a). We also found 3 mutations in bacteria with a significant negative effect on POA: a nonsynonymous mutation in *ompF* and two deletions Δ777bp in *insB* and Δ141bp in *malT*; whereas for phage we identified a nonsynonymous mutation in *J* (Fig. 4a and Supplementary Fig. S4a).

**Figure 4.**
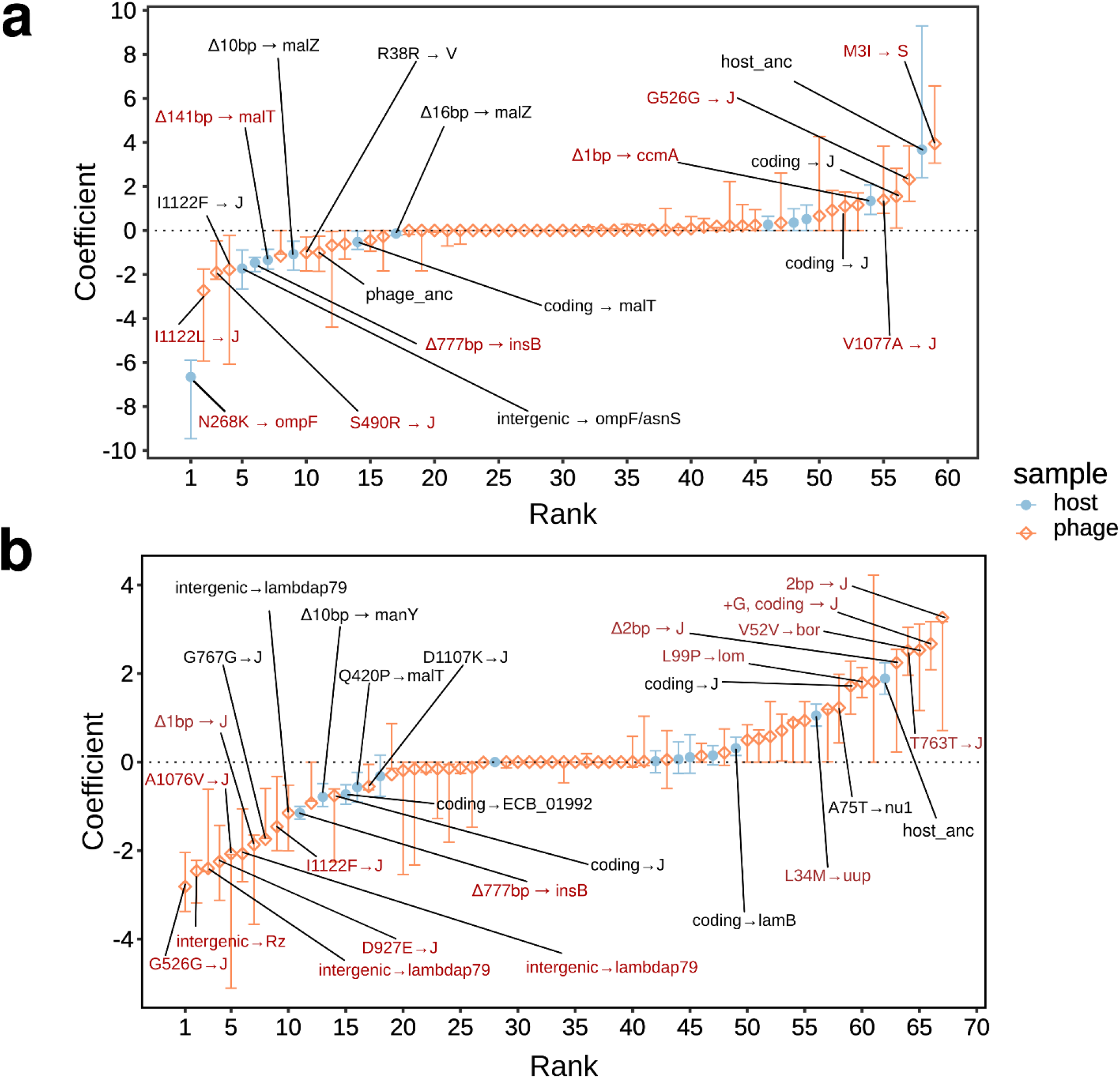
Rank ordered (most negative to most positive) coefficients inferred for putative mutations. The importance of mutational features was measured by the absolute value of the coefficients learned from each model. Error bars indicate 0.9 quantile. Labels indicate “mutation → gene” when the 90th quantile excludes 0. Mutations in red have the highest positive (negative) coefficients which lowest (highest) value is larger (smaller) or equal to 0 (from 200 bootstrap runs) and are discussed in the main text. Important features for (a) the final model predicting POA include a total of 59 non-zero coefficients, and (b) 67 non-zero coefficient values for the final model predicting EFF. The complete lists of mean, maximum and minimum values of the coefficients associated to mutations predicting POA and EFF are shown in Supplementary Data S3 and Supplementary Data S4 respectively. For further details see Methods section “Feature importance analysis”

For the EFF phenotype, 16 mutations are predicted to have a significant effect (7 positive and 9 negative; Fig. 4b, Supplementary Fig. S4b and Supplementary Data S4). Of the 7 positive predicted features, only 1 is bacterial, a nonsynonymous mutation in the *uup* gene which encodes a UvrA-like ABC family ATPase. For phage, we identified 2 insertions, 1 deletion, and 1 synonymous mutation in *J* gene, another synonymous mutation in *bor* gene, and a nonsynonymous mutation in the *lom* gene that should increase the efficiency of infection. Of the 9 features that negatively impact EFF, 1 is in the bacteria and 8 are in phage. The only bacterial mutation that negatively affects the EFF was already identified by the POA model: the Δ777bp deletion in *insB*. For the phage we identify 2 different intergenic mutations with significant negative effects downstream of *lambdap79* gene; 3 nonsynonymous, 1 synonymous (that was positive for POA) and Δ1bp deletion mutations in *J* gene and 1 intergenic mutation between *Rz* and *bor* genes (Fig. 4b, Supplementary Fig. S4b).

Results from feature importance analysis demonstrate the ability of the machine learning inference framework to identify candidate, pivotal genes involved in phage-bacteria interactions across various available phenotypes while recapitulating known biology of *λ* phage infection. We find mutations in the bacterial *malT* gene, a trans positive regulator of LamB (Debarbouille et al., 1978; Blanche et al., 2013; Maynard et al., 2010; Meyer and Lenski 2020), and several mutations located in the phage *J* gene region that were important for both POA and EFF phenotype predictions. The *J* gene encodes the tail fiber of phage *λ* which is critical to the process of adsorption to the host and injecting phage DNA via LamB (Ge and Wang 2024; Wang, Hofnung and Charbit 2000; Werts et al., 1994; Wang et al., 1998; Maddamsetti et al., 2018). Therefore, mutations in both *malT* and *J* gene region are expected to impact the phage-bacteria interaction network and the relative efficiency of infection – consistent with our model predicting the mutations to be important for both POA and EFF phenotypes. A nonsynonymous mutation in the outer membrane porin OmpF, is the most important feature for predicting a decrease in POA, but was not found to be important for predicting EFF. This mutation is shared by 10 host strains, 2 of which were sampled from day 28 and 8 were from day 37. These 10 host strains were super-resistant, that is, they were resistant to the ancestral phage *λ* strain, and all the phage isolates from the coevolution experiment. Previous studies on this bacterial population showed that phage *λ* evolves to use OmpF as a second receptor after *E. coli* evolves to down-regulate LamB (Meyer et al., 2012). Therefore, this OmpF mutation is expected to confer resistance to these evolved phage *λ* strains and so affects the POA (host-range), but not the EFF (efficiency of phage infection). Similar OmpF mutations have been described to provide resistance to a related phage, phi21, after it similarly evolved to use OmpF (Borin, Lee, Lucia-Sanz et al., 2023). Both models also identified mutations in *manY* which is an inner membrane transporter that enables phage *λ* to inject its DNA into the cytoplasm. Mutations in this protein or others in the ManXYZ complex are known to confer resistance to *λ* (Erni and Kocher 1987; Burmeister et al., 2021; Borin, Lee, Gerbino et al., 2023), and all of them impacted negatively both POA and EFF phenotypes. Most interestingly, both models were able to identify the importance of Δ777bp deletion in *insB* by an IS element from *E. coli* which affects genes not previously identified to interact with phage *λ* (Blanche et al., 2013; Maynard et al., 2010), but was recently identified to confer resistance through epistasis with other resistance mutation in *malT* through an unknown mechanism (Gupta, Peng et al., 2022).

## Discussion

In this study, we developed a machine learning framework leveraging hierarchical logistic regression to predict phage-bacteria interactions by linking infection phenotypes with genetic mutation profiles of both phage and bacterial host. By comparing models incorporating increasing layers of genotype interactions, we found that incorporating independent and additive mutational effects of phage and bacteria had the highest predictive value in inferring phenotype from genotype. In doing so, the framework identified gene regions already recognized in mediating the efficiency of infection for bacteriophage *λ* and *E. coli* (Gupta, Zaman et al., 2022; Meyer et al., 2012; Blanche et al., 2013; Burmeister et al., 2021) and predicted mutations that confer a resistant phenotype in bacteria through epistasis with other mutations (Gupta, Zaman et al., 2022). The model also identified features that were located in the phage gene *J* region, including a number of synonymous mutations as well as insertions and deletions that in principle should be detrimental, but have been shown to modulate host-range expansion and counter-defense through recombination (Borin et al. 2021). Finally, the framework identified potentially novel sites that impact phage-bacteria interaction in this bacteriophage *λ* and *E. coli* system.

Interaction inference was enabled by comparing the performance of different combinations of genotype information in predicting phage-bacteria interaction phenotypes. Model performance analysis revealed the additive model as the best predictor of interaction phenotype from phage and bacterial genotype. In the additive model, phage and bacterial mutations act independently, rather than synergistically (whether positively or negatively), to determine the infection outcome. Hence complex interaction networks may be (partially) predictable based on direct effects rather than relying on direct inference of complex interactive effects that are more challenging to measure (Tamar and Kishony 2022). This result may be limited by sampling and does not exclude the possibility that higher order gene-gene interactions affect infection phenotypes. The number of phage-host mutation pairs scales as the product of the number of phage and host mutations in higher order models (including both the nonlinear and mixed models), but most of these mutational combinations were not observed in our set of sequenced strains. As such, fitting higher order models leads to underdetermined systems even with the introduction of regularization terms meant to limit the number of weak contributions from mutations – whether direct or in combination. Future work would have to significantly scale-up genotyped combinations of overlapping mutations in different contexts to robustly infer phage-bacteria interaction mutational pairs.

The machine learning inference framework was able to detect the importance of previously identified adaptive mutations that modify phage-host interactions and potentially novel genes and mutations that modulate qualitative and quantitative features of virus-microbe interactions. Although identification of novel genes is not expected to be comprehensive, we did identify several nonsynonymous mutations that impact POA and EFF phenotypes and that are not directly involved in phage-host surface recognition – notably, mutations in *uup*, and *ccmA* genes in bacteria and *S* and *lom* genes in phage. The nonsynonymous mutation in the phage *S* gene region is identified as important for predicting the presence (or absence) of infection. This gene encodes a holin – a small inner membrane protein required for phage-induced host lysis (Chang, Nam and Young 1995). Thus, we interpret the feature analysis to imply that a mutation in the S gene has a direct impact on the lysis of host cells and resulting POA/EFF phenotypes. Similar mutations were uncovered via experimental evolution to counteract a gene deletion in the host that helps facilitate phage DNA replication (Gupta et al., 2020). This mutation may potentially extend the infection process and allow phage more time to initiate DNA replication in the debilitated host, increasing the chance of a successful infection. Another candidate mutation that shapes EFF interaction phenotypes was found in the phage *lom* gene region. We note that this site was previously reported to increase phage resistance through an unknown mechanism (Borin et al., 2021), future work may prioritize the impact of mutations in this gene region on within-cell viral dynamics. Finally, we note that despite the potential for false positives and negatives, evolutionary effects including genetic hitchhiking and recombination may move adaptive mutations onto different backgrounds, improving detection of driver mutations of infection.

In summary, we have developed a machine learning framework for predicting genotypic drivers of both the qualitative and quantitative nature of host-pathogen interactions that can serve as a foundation for similar analyses in other co-evolutionary contexts where strong selection pressures enable the selection of virus and host mutations that modulate interaction phenotypes. Moving forward, it will be essential to investigate how variations in host-phage systems, ecological contexts, and genetic constraints introduce sufficient diversity during coevolution to allow phenotype prediction from genotype and we acknowledge that determining a minimum viable training sample size for consistent model performance would be a valuable direction in future applications. Nevertheless, this framework could help prioritize research on identifying novel drivers of infection, focusing efforts on mutations most likely to alter phage-bacteria phenotypes. Although we applied this framework in a relatively low genetic diversity context, this data-driven approach does not require *a priori* knowledge of driver genes and mutations. Hence, this inference framework could be applied to other, even poorly characterized, phage-bacteria systems, potentially improving understanding of interactions in complex, natural systems as well as for phages that target bacterial pathogens.

## Materials and Methods

### Experimental setup and data collection

We analyzed phenotypes and genomes associated with *E. coli* B strain REL606 and phage *λ* strain cI26, cocultured for a 37-day period. Samples were taken on checkpoint days for pairwise quantitative plaque assays as described in (Gupta, Peng et al., 2022). The EOP value measures the efficiency of a phage infecting a derived host strain relative to that for infecting the ancestral strain. The EOP value for a phage, *j*, infecting a host, *i*, is computed as:

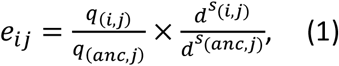

where *q*_(*i,j*)_ is the number of plaques for phage *j* against host *i, q*_(*anc,j*)_ is the number of plaques for phage *j* against the ancestral host strain, *s*_(*i,j*)_ is the number of dilutions performed to observe distinguishable and countable clear plaques for phage *j* against host *i, s*_(*anc,j*)_ is the number of dilutions performed to observe distinguishable and countable clear plaques for phage *j* against the ancestral host strain and *d* is the dilution ratio which is 5 in our experiment. A positive EOP value from the cross-infection plaque assay indicates a successful infection event for a given phage-host pair. In contrast, a zero EOP value indicates the phage has no capacity to infect. A larger EOP value from the cross-infection plaque assay indicates that the phage can infect a given host more efficiently than the ancestral host strain.

For each phage and host samples taken from each checkpoint, the DNA extraction, library preparation and sequencing experiment was performed as described in (Gupta, Peng et al., 2022). Mutation profiles based on the genome sequencing data were constructed using *breseq* as described in (Gupta, Peng et al., 2022). In addition to the mutations revealed by *breseq* results, for both host and phage we created an artificial mutation as the indicator for the ancestral strain to add the ancestral strain into the mutation profile table. For this artificial mutation, only the ancestral strain is indicated to have this mutation. All other strains were shown to not have this mutation in the mutation profile table.

### Feature construction

For a total number of *U* host samples and *V*phage samples, we denote the EOP value for the *i*-th host against *j*-th phage as *e*_*ij*_ where *i* ∈ [1, *U*] and *j* ∈ [1, *V*]. Let *N* be the total number of unique mutations observed for the host and *M* be the total number of unique mutations observed for the phage, the host mutation profile *H* is a matrix of dimension *U* by *N*, and the phage mutation profile *P* is a matrix of dimension *V* by *M*. Let *h*_*il*_ be an element from *H*, then *h*_*il*_ = 1 corresponds to the presence of the *l*-th mutation in the *i*-th host whereas *h*_*il*_ = 0 corresponds to the absence of the *l*-th mutation in the *i*th host. Similarly, let *p*_*jk*_ be an element from *P*, then *p*_*jk*_ = 1 corresponds to the presence of the *k*-th mutation in *j*-th phage whereas *p*_*jk*_ = 0 corresponds to the absence of the *k*-th mutation in the *j*-th phage.

Five sets of features were constructed based on the mutation profiles of the host and phage. The H-only model is constructed based on a linear combination of ‘host only’ mutation profiles. The H-only model, denoted as 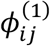, can be represented as:

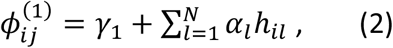

where *γ*_1_ represents a scalar of the bias term and *α*_*l*_ is the coefficient for the *l*-th host mutation. *γ*_1_ and *α*_*l*_ will be learned from the model. The H-only model can also be represented in matrix form as:

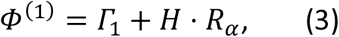

where *Γ*_1_ is a *U* by *V* matrix by repeating *γ*_1_, i.e. *Γ*_1_ = [*γ*_1_]_*U*×*V*_, *R*_*α*_ is a *N* by *V* matrix by stacking the same coefficient vector *α* horizontally, i.e. [*α*|*α*| … |*α*|*α*]_*N*×*V*_.

The P-only model is constructed based on a linear combination of ‘phage only’ mutational profiles. The P-only model, denoted as 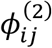, can be represented as:

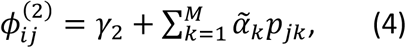

where *γ*_2_ represents a scalar of the bias term and 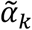 is the coefficient for the *k*-th phage mutation. *γ*_2_ and 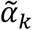 will be learned from the model. This model can also be represented in matrix form as:

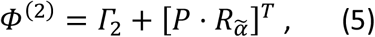

where *Γ*_2_ is a *U* by *V* matrix by repeating *γ*_2_ and 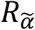 is a *M* by *U* matrix by stacking the same coefficient vector 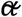 horizontally, i.e. 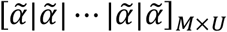.

The linear model, denoted as 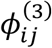,utilizes a linear combination of phage and host mutational features and can be represented as:

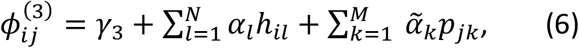

where *γ*_3_ represents a scalar of the bias term, *α*_*l*_ is the coefficient for the *l*-th host mutation and

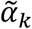 is the coefficient for the *k*-th phage mutation. *γ*_3_, *α*_*l*_ and 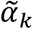 will be learned from the model. The linear model can also be represented in matrix form as:

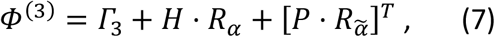

where *Γ*_3_ is a *U* by *V* matrix by repeating *γ*_3_, i.e. *Γ*_3_ = [*γ*_3_]_*U*×*V*_, *R*_*α*_ is a *N* by *V* matrix by stacking the same coefficient vector *α* horizontally, i.e. [*α*|*α*| … |*α*|*α*]_*N*×*V*_ and 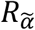 is a *M* by *U* matrix by stacking the same coefficient vector 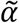 horizontally, i.e. 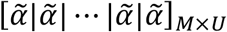 The assumption for the linear model is that the impact of mutations from both the phage and host have additive effects on the observed outcome.

The nonlinear model, denoted as 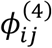, utilizes nonlinear combination of phage and host mutational features as the input and can be represented as:

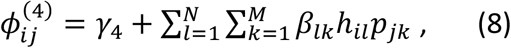

where *γ*_4_ represents a scalar of the bias term, *β*_*lk*_ denotes the coefficient for the *l*-th host mutation and *k*-th phage mutation in the corresponding *i*-th host and *j*-th phage pair. *γ*_4_ and *β*_*lk*_ will be learned from the model. This nonlinear model can also be represented in the matrix form as:

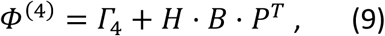

where *Γ*_4_ is a *U* by *V* matrix by repeating *γ*_4_, i.e. *Γ*_4_ = [*γ*_4_]_*U*×*V*_, *B* is the *N* by *M* coefficient matrix. The assumption for the nonlinear model is that the impact of the genetic mutations on the observed outcome comes from the additive effects of co-occurring phage-host mutation pairs. In other words, *h*_*il*_*p*_*jk*_ = 1 only when both the host *i* has mutation *l* and phage *j* has mutation *k*.

Based on the formulation of the linear and nonlinear models, it is natural to combine both effects to get a more sophisticated input feature, by adding up both effects. The mixed model, denoted as 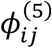, utilizes a mixed combination of linear and nonlinear effects of host and phage mutation features and can be represented as:

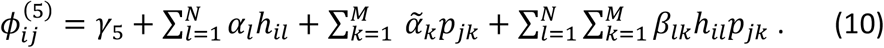

The matrix form of the mixed model is:

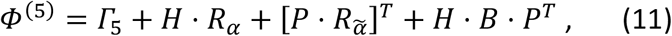

where *Γ*_5_ is a *U* by *V* matrix by repeating *γ*_5_, i.e. *Γ*_5_ = [*γ*_5_]_*U*×*V*_.

### Framework design

We designed a framework comprised of two types of predictions. First, we designed a framework that predicts the phage-host cross interaction network (i.e., host range). This model tries to find the set of mutational features that best distinguish between presence (EOP > 0) and absence (EOP = 0) of infection (POA) using classification models. Second, we built a framework to predict the efficiency of the interaction (EFF). This model is designed to find the set of mutational features that best predicts the efficiency of a phage infecting a host in those phage-host pairs where the host is susceptible to the phage (EOP > 0).

### Model for predicting phage-host cross-interaction network (POA)

In order to determine the presence or absence of a successful infection event for a phage-host pair, we binarized the EOP values *e*_*ij*_ into 0 and 1, i.e.

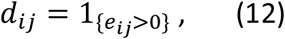

where *d*_*ij*_ = 0 indicates a failure of the infection and *d*_*ij*_ = 1 indicates success. Here we used logistic regression to model the relationship between mutation profiles and the existence of successful infection in phage-host pairs, that is:

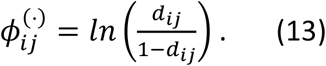

Each of the five sets of features, namely H-only, P-only, linear, nonlinear and mixed, were used as the input features for the models 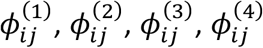 and 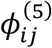 respectively. In practice, we used LASSO for feature selection and regularization. The penalty term parameter for LASSO was determined by using 10-fold cross-validation on the training data. The prediction classification error, 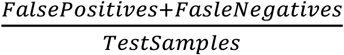, was used to assess the performance for this model. The mean classification error was calculated by taking the mean of classification error from 200 runs.

### Model for predicting the efficiency of phage-bacteria infections (EFF)

We applied a log transformation on the positive EOP values to normalize the distribution (Supplementary Fig. S5a). For a given phage-host pair where a successful infection event is present, that is *e*_*ij*_ > 0, we denote the natural log transformed EOP value as:

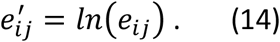

Shapiro-Wilk test was performed to check the normality of the distribution of 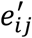 (Supplementary Fig. S5b).

Linear regression was used to model the relationship between mutation profiles and the intensity of successful infections in phage-host pairs, that is:

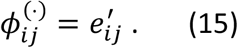

Each of the five sets of features, namely H-only, P-only, linear, nonlinear and mixed, were used as the input features for the models 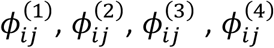 and 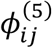 respectively. For the linear model, we also used LASSO for feature selection and regularization. The penalty term parameter for LASSO was determined by using 10-fold cross-validation on the training data. Finally, the MAE was used to evaluate the performance of the model.

### Train-validation split and feature evaluation

To assess the performance of different features for the logistic regression model, we performed 200 bootstrap runs to predict the existence of phage infection. Specifically, in each run the training set was generated by randomly select *U* × *V* samples from the entire dataset with replacement. The *d*_*ij*_ values that were not selected as training samples form the validation set. As a control, for each run, a null model was built to predict the outcomes by randomly sample *d*_*ij*_ values from a Bernoulli distribution *Bern* 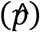 where 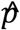 is the maximum likelihood estimator (MLE) of the proportion of successful infection from the training set of that run. After the 200 runs, the training and validation prediction error were compared between pairs of the models including the null model and models based on phage and host mutations only and linear, nonlinear, and mixed combinations of phage and host mutational features.

Similarly, we also performed 200 bootstrap runs for the linear model to predict the infection efficiency. Specifically, in each run the training set was generated by randomly sample 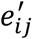 with replacement. The size of 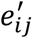 sampled as the training set in each run matches the total number of the 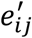 The 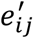 that were not selected in the training set forms the validation set. As a control, for each run, a null model was built by always predicting the efficiency of infection as the mean 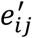 of the training set for that run. After the 200 runs, the training and validation MAEs were compared between pairs of the models including the null model and every feature model set.

### Final predictions and feature importance analysis

After comparing the training and validation performance of models based on the different mutational sets with 200 bootstrap runs, a final model, that integrates predictions of POA and EFF was constructed. The penalty term parameter for each of the prediction frameworks was chosen as the mean of the best penalty term parameter from each of the 200 bootstrap runs. After model fitting, the predicted outcome 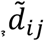 for the POA model and 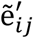 for the EFF model was calculated. For each step of the final models, the importance of feature was measured by the absolute value of coefficients learned from each step.

## Author Contributions

Conceptualization: JSW, CYL & JRM

Methodology: ALS, SP, CYL & JSW

Investigation: ALS, SP, CYL, AG

Visualization: ALS, SP

Writing – original draft: ALS, SP & JSW

Writing – review & editing: ALS, SP, CYL, AG, JRM, JSW

## Software Availability

https://github.com/aluciasanz/genotype_to_phenotype_inference_model

## Acknowledgments

JSW - Army Research Office (W911NF1910384), NSF (2200269), NIH (R01 AI146592), Simons

Foundation (930283), Chaires Blaise Pascal of the Île-de-France region.

JRM - Howard Hughes Medical Institute Emerging Pathogens Initiative grant 311169.

