## Supplementary for "Inferring strain-level mutational drivers of phage-bacteria interaction phenotypes arising during coevolutionary dynamics"

Lucia-Sanz, et al.

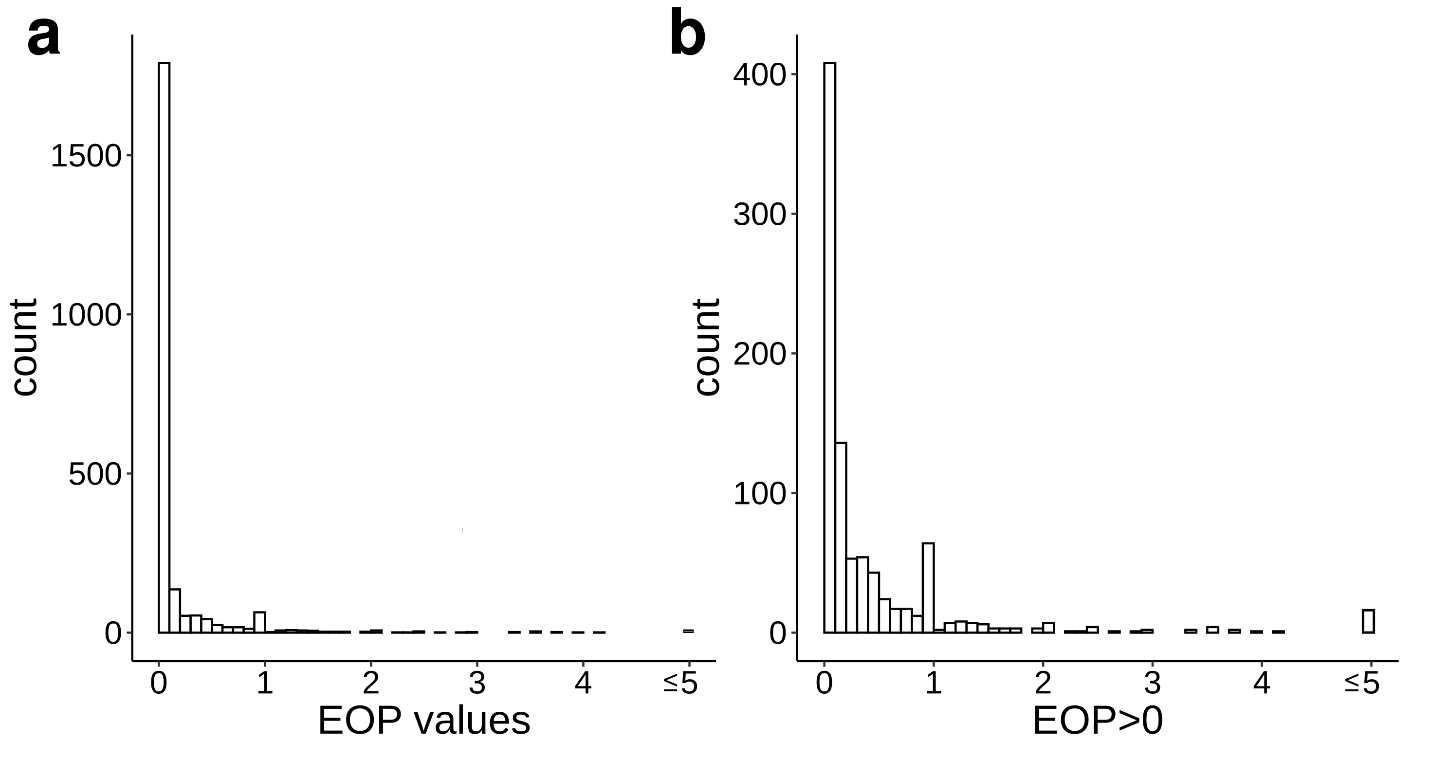

**Supplementary Figure S1. Distribution of the experimentally obtained EOP values.** (a) Original distribution of the EOP values for 2295 phage-host infection pairs. (b) Distribution of 913 positive EOP values. Bin width=0.1.

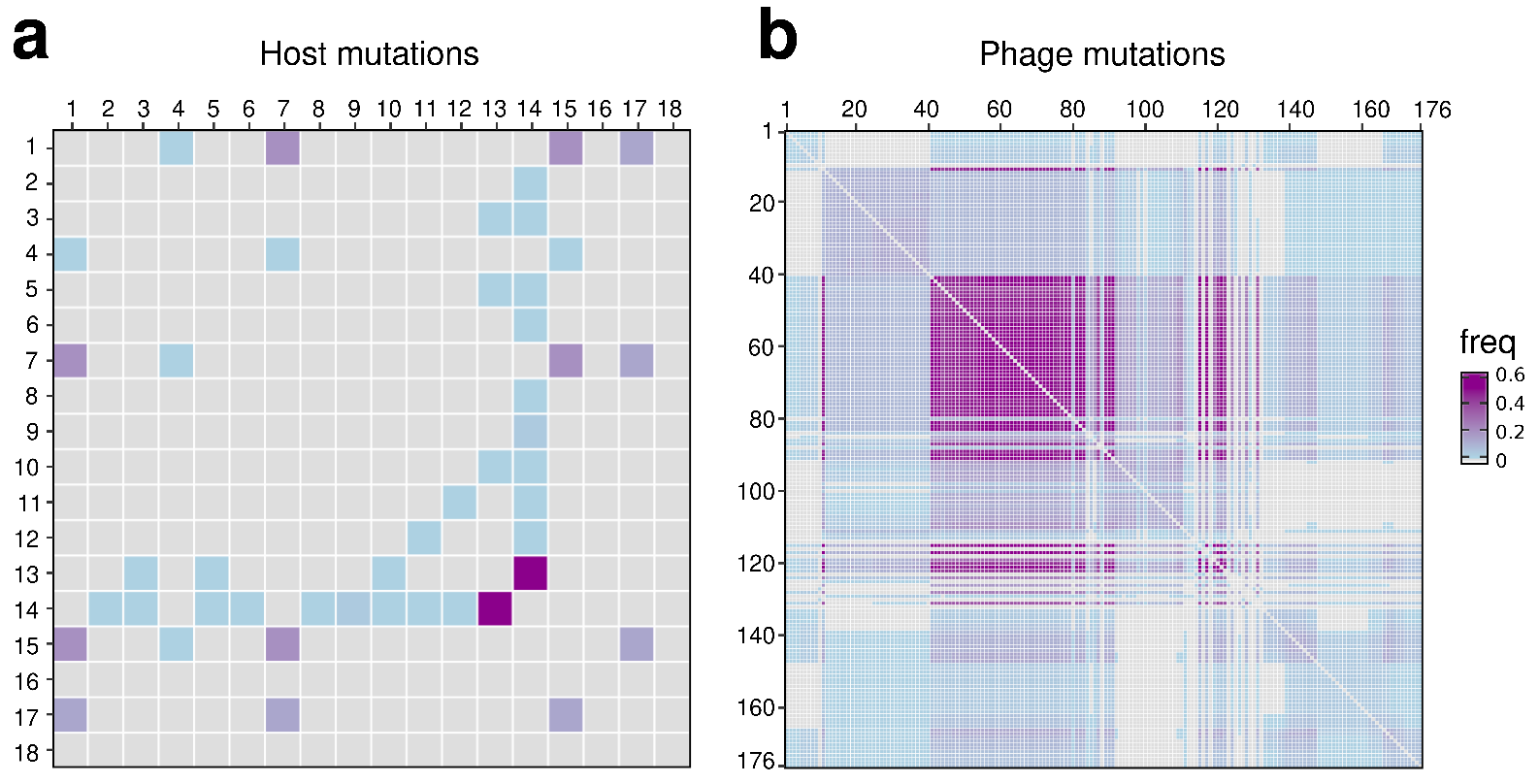

**Supplementary Figure S2. Correlations of mutational appearances in host and phage.** (a) 18x18 host and (b) 176x176 phage mutation matrices representing the frequency with which pairs of mutations simultaneously appear within the same genetic background.

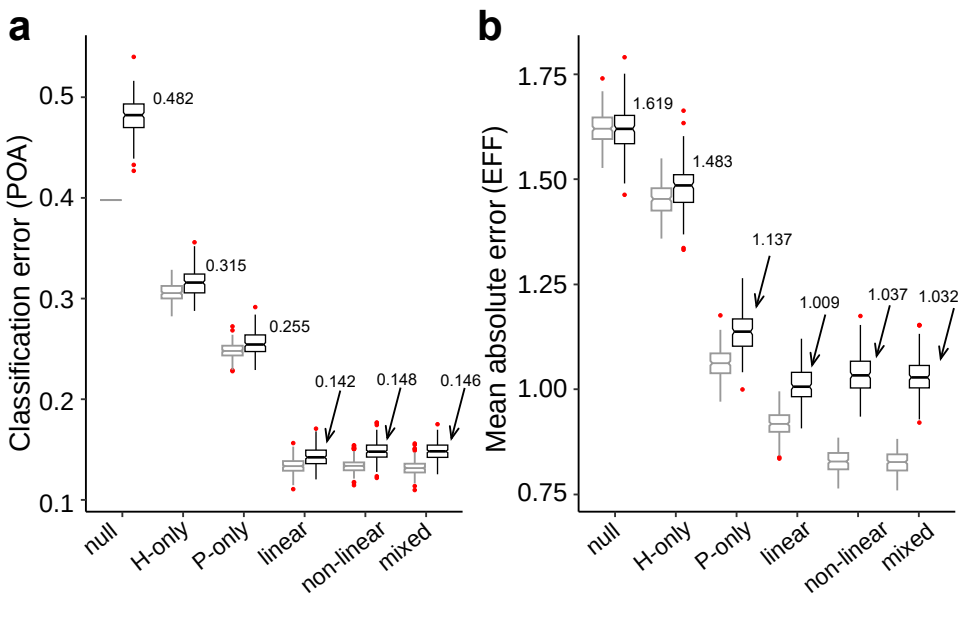

**Supplementary Figure S3: Model performances for different feature sets.** The lowest mean value in the validation set for POA and EFF models corresponds to the linear model. (a) Classification error distributions in the training (grey) and validation (black) sets for the predictions of the phage-host interaction network (POA) (ANOVA post hoc Tukey p<0.01). The lowest mean value in the validation set corresponds to the linear model (classification error = 0.142). (b) Mean absolute error distributions in the training (grey) and validation (black) sets for the predictions of efficiency of infection (EFF) (ANOVA post hoc Tukey p<0.001, comparing different mutation feature models and a null model. The lowest mean value in the validation set corresponds to the linear model (MAE = 1.009). Boxplots contain 25^th^-75^th^ percentiles, whiskers indicate minimum and maximum values, middle lines are the median (value indicated) of 200 bootstrap runs. Red dots are outliers. For more details see Method section “Train-validation split and feature evaluation”

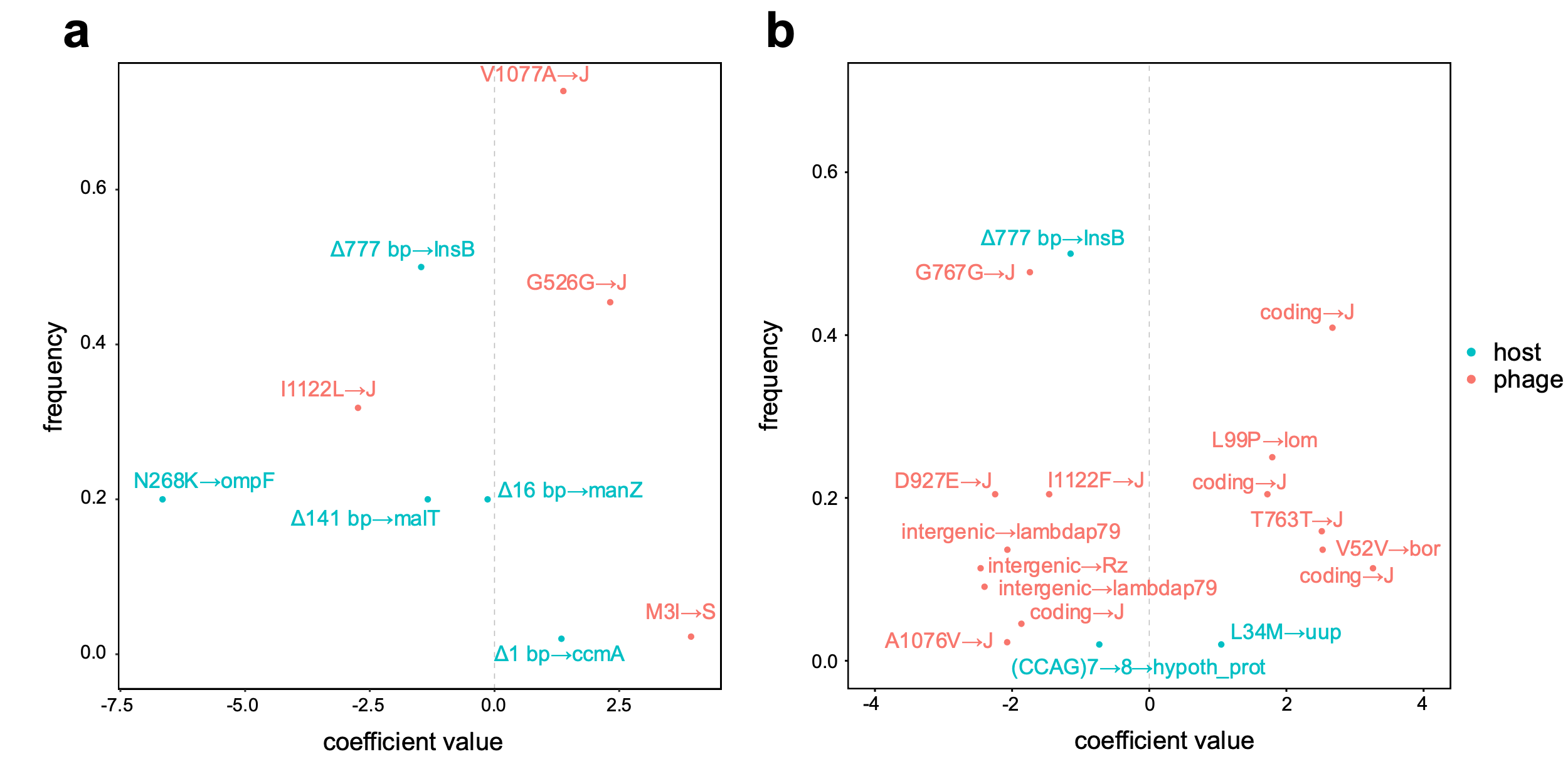

**Supplementary Figure S4. Frequency and inferred coefficients of putative important mutations.** Comparison of the observed frequencies of phage (orange) and host (blue) mutations in the dataset with the inferred coefficients resulted from feature importance analysis of the linear model predicting (a) POA and (b) EFF phenotypes.

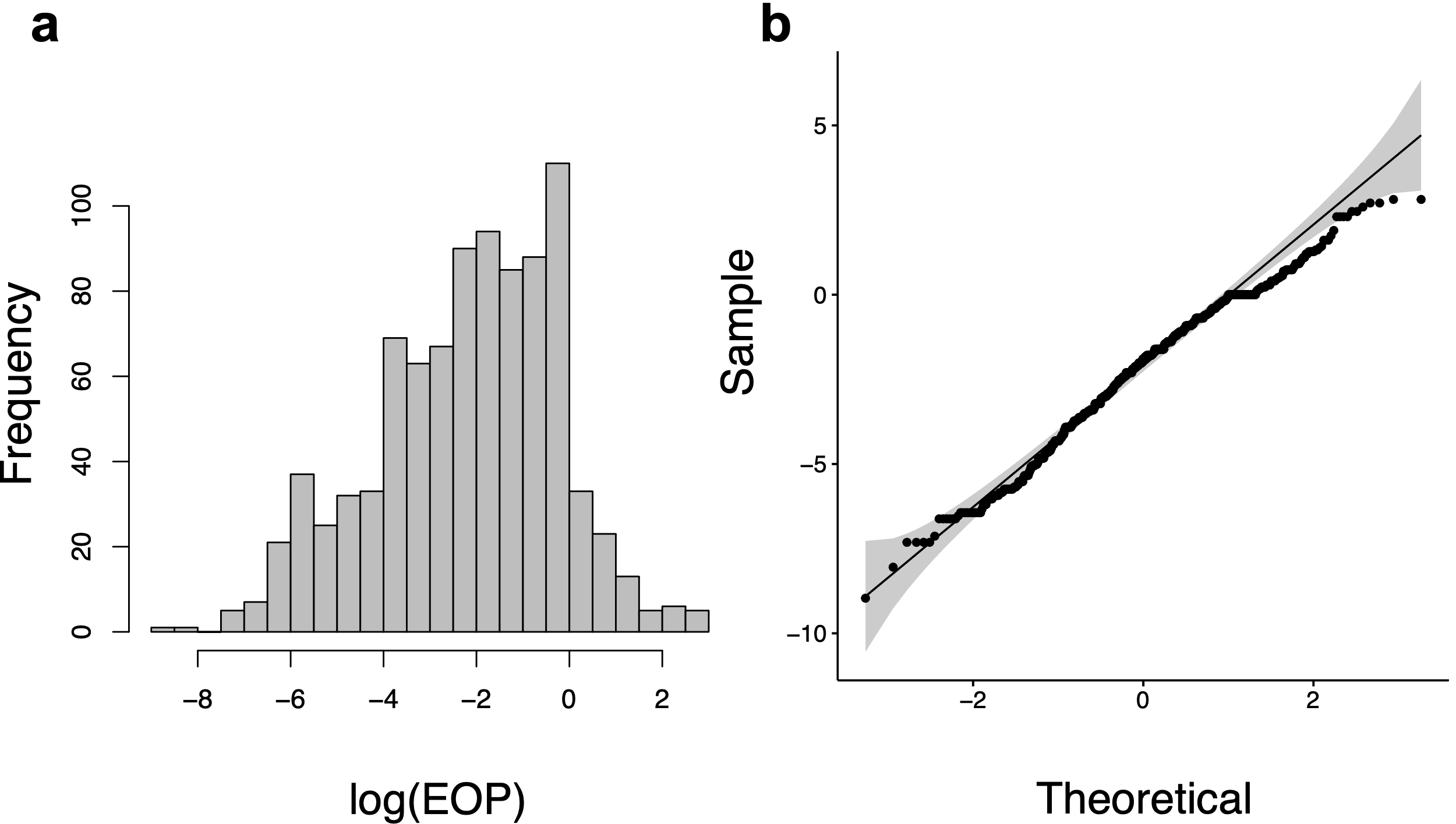

**Supplementary Figure S5. Log transformed positive EOP value distribution.** (a) Distribution of the log positive EOP values (b) Q-Q plot for log positive EOP values against normal quantiles (Shapiro-Wilk test *P* value = 3.283e-8).

**Supplementary Data S1. Mutation profile tables for host and phage.**

**Supplementary Data S2. Ordered features with non-zero coefficients from final model for predicting POA based on a linear combination of phage and host mutation profiles.**

**Supplementary Data S3. Ordered features with non-zero coefficients from final model for predicting EFF based on a linear combination of phage and host mutation profiles.**
